# RWRDC: Predicting Efficacious Drug Combinations in Cancer Based on Random Walk with Restart

**DOI:** 10.1101/2020.09.13.295063

**Authors:** Qi Wang, Guiying Yan

## Abstract

**Background:** Compared with monotherapy, efficacious drug combinations can increase the therapeutic effect, decrease drug resistance of experimental subjects and the side effects of drugs. Therefore, efficacious drug combinations are widely used in the treatment of complex diseases, such as various cancers. However, compared with the mathematical model and computational method, experimental screening efficacious drug combinations is time-consuming, costly, laborious, and inefficient;

**Methods:** we predicted efficacious drug combinations in cancer based on random walk with restart (RWRDC). An efficacious score can be obtained between any two individual drugs by RWRDC;

**Results:** As a result, we analyzed the rationality of the efficacious score first. Besides, compared with the other methods by leave-one-out cross-validation, all the Area Under Receiver Operating Characteristic Curves (AUROCs) of RWRDC were higher for data sets of breast cancer, colorectal cancer, and lung cancer. Moreover, the case study of breast cancer showed that RWRDC could discover potential efficacious drug combinations;

**Conclusions:** These results suggest that RWRDC is a novel way to discover efficacious drug combinations in cancer, which provides new prospects for cancer treatment. Furthermore, RWRDC is a semi-supervised learning framework that can be used to predict combinations of drugs for other complex diseases.

## I. Background

A combinatorial drug is a fixed-dose combination, including two or more active drug components in a single dosage form, which is manufactured and distributed in a fixed dose (Collier, 2012). Drug combinations are classified as efficacious or non-efficacious depending on whether they can produce expected improvements in clinical trials or preclinical studies (Liu *et al.,* 2014). Compared with monotherapy, efficacious drug combination can increase the efficacy of the therapeutic effect, and decrease the drug side-effects and drug resistance of experimental subjects. Especially, drug combinations interact with multiple targets in the molecular networks of complex diseases, as a consequence, efficacious drug combinations is targeted treating a large number of complex diseases, such as various cancers (Feliu *et al.,* 2009), human immunodeficiency virus (HIV) as well as fungal (Groll and Tragiannidis, 2009) and bacterial infections (Tamma *et al.,* 2012), etc.

Although efficacious drug combinations are important and meaningful, there are too many plausible combinations of individual drugs, and it is time-consuming and costly to screen efficacious drug combinations by experiment. Mathematical models and computational methods show their advantages in drug screening. For example, Kristina Preuer *et al.* (Preuer *et al.,* 2018) presented a novel deep learning model, DeepSynergy, to predict synergy scores of drug combinations for cancer cell lines. They used chemical and genomic information as model inputs and took into account standardization strategies for heterogeneity of input data. Janizek *et al.* (Janizek *et al.,* 2018) used the XGBoost method combing with a tool named TreeSHAP to predict drug combinations and make prediction results explainable. Zhang *et al.* (Tianyu Zhang *et al*, 2018) proposed a novel deep learning model to predict drug combinations by integrating multi-type of genomics data and chemical structure data. They uncovered potential mechanisms of the synergy of specific drug combinations against given cancer cell lines. Chen *et al.* (Chen *et al.,* 2016) developed a model named Network-based Laplacian regularized Least Square Synergistic drug combination prediction (NLLSS) to predict potential synergistic drug combinations. They also carried out biological experiments, finally, 7 of the 13 antifungal synergistic drug combinations predicted by the NLLSS were verified. (Bai *et al.,* 2018) devised a novel improved naïve Bayesian algorithm to construct classification models to predict effective drug combinations. (Xu *et al.,* 2017) predict effective drug combinations by integrating biological, chemical and pharmacological information based on a stochastic gradient boosting algorithm. (Li *et al.,* 2020) used neighbor recommender method combined with ensemble learning algorithms to predicting drug combinations, and they predicted seven candidate drug combinations for paclitaxel, and successfully verified the promising effects of two of the predicted combinations.

Most computational models about predicting efficacious drug combinations are using supervised learning methods. In fact, due to the very few known non-efficacious drug combinations, it has been very difficult to obtain all the label information (efficacious or non-efficacious) of the training set. In this paper we used a semi-supervised learning method, called random walk with restart model, to predict efficacious drug combinations in three specific cancers (RWRDC). The random walk is a powerful mathematical model that is evolving rapidly in some important tasks, such as ranking (Minkov and Cohen, 2007; Bubnicki and Orski, 2005), similarity and recommendation (Li *et al.,* 2012; Tayebi *et al.,* 2011), or link prediction (Backstrom and Leskovec, 2011). Then, we analyzed the value of restart probability and the rationality of scoring for RWRDC. Moreover, our model achieved reliable predictions with cross-validation and it obtained higher the Area Under Receiver Operating Characteristic Curves (AUROCs) than other methods for three data sets. A case study of breast cancer demonstrated the effectiveness and potential of RWRDC in identifying potential drug combinations. Finally, we discussed the strengths and limitations of RWRDC and its prospects.

## II. Results

In this section, we first analyzed the value of restart probability and the rationality of scoring for RWRDC. Next, cross-validation evaluated the predictive performance of RWRDC. Then, we conducted a case study to verify the efficiency of RWRDC in discovering novel combinatorial drugs.

### A. Effects of restart probability

A receiver operating characteristic (ROC) curve is a graphical plot that shows the diagnostic ability of the binary classifier system because its recognition thresholds are different (Fawcett, 2006). AUROC (Area Under Receiver Operating Characteristic Curve) is the area under the ROC curve with a value between 0 and 1. AUROC can intuitively evaluate the quality of classifier, the larger the value, the better.

Leave-one-out cross-validation (LOOCV) is a widely used cross-validation, especially when the data size is not very large. specifically, one positive sample (a known efficacious drug combination) was taken as the test set in turn, the other positive samples (other known efficacious drug combinations) were taken as the training set, and the other non-positive samples (other possible drug combinations) were taken as the *CS*. After scoring by RWRDC, every sample of *PS* and *CS* was scored and sorted. After taking all positive samples as the test set in turn, the score of the test set was compared with the scores of all candidate samples, then the predictive performance of the RWRDC was evaluated by changing the preset threshold.

To determine the effect of restart probability 1 — *c* on the performance of RWRDC, we calculated AUROCs with various values of 1 — *c* ranging from 0.05 to 0.95 by LOOCV. **Figure 1** shows the performance of different restart probabilities on the LOOCV in the different data sets. The orange line represented the average AUROC when the restart probability changed. It could be observed that the performance of RWRDC was better when restart probability was about 0.05. Therefore, we set 1 — *c* to 0.05.

**Fig. 1.**
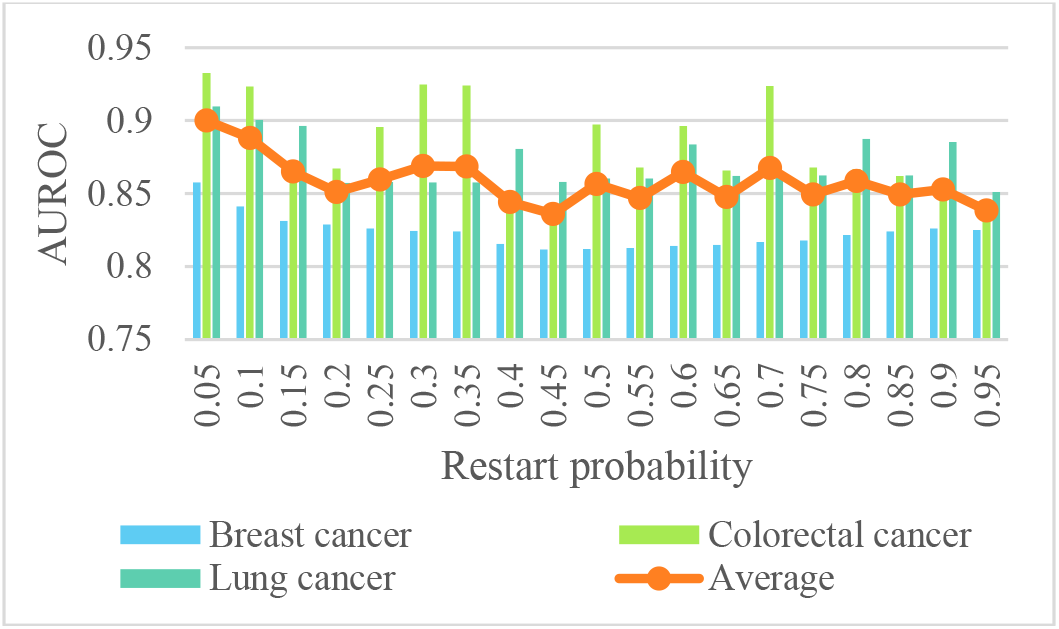
The performance of restart probability on the LOOCV.

### B. Rationality Analysis of Scoring

To check whether the drug-drug pairs with high efficacious scores were more likely to be efficaciously combined, all candidate drug-drug pairs in three data sets were ranked by RWRDC. In this paper, the average efficacious score of *PS* (efficacious combinations), *CS* (unknown combinations), and *NS* (non-efficacious combinations in *CS*) were calculated respectively. If the average efficacious score of *PS* was higher than that of *CS* and also the average efficacious score of *CS* higher than that of *NS* on different data sets, the scoring process was reasonable.

In **Table 1**, The average efficacious score of *PS, CS*, and *NS* were expressed by *PS score, CS score*, and *NS score*, respectively. As shown in **Table 1**, all efficacious scores of *NS* were close to zero, drug-drug pairs with higher efficacious scores were more likely to be efficacious drug combinations of three data sets.

**TABLE I.** Comparing the efficacious scores on different data sets

| Data sets | PS score | CS score | NS score |
| --- | --- | --- | --- |
| Breast cancer | 0.0425 | 1.5589e-4 | 6.9571e-5 |
| Colorectal cancer | 0.0659 | 2.2667e-4 | 5.9617e-4 |
| Lung cancer | 0.0519 | 3.1624e-4 | 1.8237e-7 |

### C. Cross-validation tests

In this paper, we used LOOCV for different methods and data sets to test the predictive ability of the model. Briefly speaking, the drug-drug pairs corresponding to the elements of 1 in the adjacent matrix A were proportionally used as the training set and the test set, and the elements of 0 in the adjacent matrix A were used as the *CS*. Then the efficacious score of each sample of the test set and *CS* was predicted.

Besides, the model was compared with the random walk model and NLLSS for different data sets. To compare the predictive power of different models more fairly, we used the same data, that is, we only used the data of existing efficacious drug combinations, and we did not use data such as the similarity between drugs. Especially, in the NLLSS model, we regarded each drug as the main drug and the adjuvant drug.

As shown in **Figure 2**, in the case of LOOCV, the blue line represented the ROC curve of RWRDC, the red line and orange line represented the ROC curve of the random walk model and NLLSS, respectively. The graphs show that the prediction result of RWRDC is better than that of the random walk model and NLLSS.

**Fig. 2.**
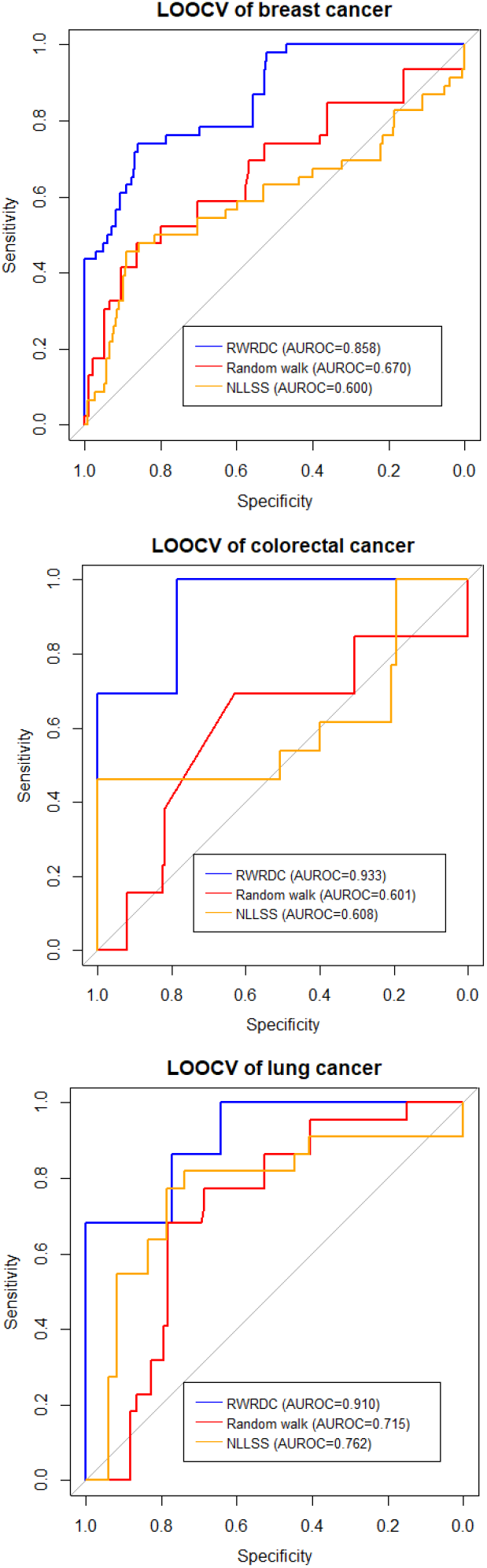
The ROC curves of breast cancer, colorectal cancer and lung cancer with LOOCV.

### D. Case study

The case study aimed to examine the capability of RWRDC in discovering novel efficacious drug combinations. For breast cancer, we ranked all drug-drug pairs based on their corresponding predicted efficacious scores. Prediction results were verified based on not only the DCDB database but recently published experimental literature. The efficacious drug combinations account for about 5%-9% of the total combinations of each cancer data set (**Table 3**), therefore, we analyzed the top 5% and 10% predictions. **Table 2** listed all the corresponding top 10% ranking results. The “Null” in the table indicates that documentation has not been found.

**TABLE II.** Case study of breast cancer

| Rank | Drug1 | Drug2 | Evidence |
| --- | --- | --- | --- |
| 1 | Arzoxifene | LG100268 | DCDB |
| 2 | Decitabine | Depsipeptide | DCDB |
| 3 | Taxane | Gefitinib | DCDB |
| 4 | PD98059 | Gefitinib | DCDB |
| 5 | Fulvestrant | Tipifarnib | DCDB |
| 6 | Olaparib | Cediranib | DCDB |
| 7 | Vinorelbine | Neratinib | DCDB |
| 8 | Cisplatin | Gemcitabine | DCDB |
| 9 | Doxorubicin | Mitomycin | DCDB |
| 10 | Doxorubicin | Metformin | DCDB |
| 11 | Methylseleninic acid | Tamoxifen | DCDB |
| 12 | Cyclophosphamide | Pemetrexed | DCDB |
| 13 | Sorafenib | Capecitabine | DCDB |
| 14 | Ixabepilone | Capecitabine | DCDB |
| 15 | Paclitaxel | Tubacin | DCDB |
| 16 | Paclitaxel | Trabectedin | DCDB |
| 17 | Paclitaxel | Toremifene | DCDB |
| 18 | Epirubicin | Trastuzumab | DCDB |
| 19 | Trastuzumab | Bortezomib | DCDB |
| 20 | Carboplatin | Trastuzumab | DCDB |
| 21 | Fulvestrant | Anastrozole | DCDB |
| 22 | Anastrozole | Goserelin acetate | DCDB |
| 23 | Everolimus | Exemestane | DCDB |
| 24 | Everolimus | Letrozole | DCDB |
| 25 | Tamoxifen | Exemestane | DCDB |
| 26 | Tamoxifen | Goserelin acetate | DCDB |
| 27 | Lapatinib | Letrozole | DCDB |
| 28 | Paclitaxel | Olaparib | DCDB |
| 29 | Vinorelbine | Trastuzumab | DCDB |
| 30 | Docetaxel | Gemcitabine | DCDB |
| 31 | Bevacizumab | Lapatinib | DCDB |
| 32 | Doxorubicin | Docetaxel | DCDB |
| 33 | Gemcitabine | Trastuzumab | DCDB |
| 34 | Lapatinib | Capecitabine | DCDB |
| 35 | Bevacizumab | Capecitabine | DCDB |
| 36 | Cyclophosphamide | Capecitabine | DCDB |
| 37 | Docetaxel | Capecitabine | DCDB |
| 38 | Paclitaxel | Bevacizumab | DCDB |
| 39 | Paclitaxel | Lapatinib | DCDB |
| 40 | Paclitaxel | Cyclophosphamide | DCDB |
| 41 | Docetaxel | Trastuzumab | DCDB |
| 42 | Doxorubicin | Trastuzumab | DCDB |
| 43 | Cyclophosphamide | Trastuzumab | DCDB |
| 44 | Bevacizumab | Trastuzumab | DCDB |
| 45 | Trastuzumab | Tamoxifen | DCDB |
| 46 | Paclitaxel | Trastuzumab | DCDB |
| 47 | PD98059 | Taxane | Null |
| 48 | Tipifarnib | Anastrozole | Null |
| 49 | Paclitaxel | Cediranib | Null |
| 50 | Neratinib | Trastuzumab | (Canonici *et al.*, 2013) |
| 51 | Fulvestrant | Goserelin acetate | (G. Steger *et al.*, 2005) |
| 52 | Letrozole | Exemestane | Null |
| 53 | Metformin | Mitomycin | Null |
| 54 | Cisplatin | Docetaxel | (Wang, 2007) |
| 55 | Everolimus | Tamoxifen | (Bachelot *et al.*, 2012) |
| 56 | Anastrozole | Tamoxifen | Null |
| 57 | Methylseleninic acid | Exemestane | Null |
| 58 | Methylseleninic acid | Goserelin acetate | Null |
| 59 | Everolimus | Lapatinib | Null |
| 60 | Cisplatin | Trastuzumab | (Pegram and Slamon, 1999) |
| 61 | Ixabepilone | Sorafenib | Null |
| 62 | Docetaxel | Mitomycin | Null |
| 63 | Metformin | Docetaxel | (Yi Z, Qingli Z, Ji M, 2018) |
| 64 | Pemetrexed | Capecitabine | (Xiaorong Y, Lingru M, Huanxin Y, et al., 2015) |
| 65 | Paclitaxel | Pemetrexed | (Tas *et al.*, 2008) |
| 66 | Trastuzumab | Mitomycin | Null |
| 67 | Metformin | Trastuzumab | (Thor *et al.*) |
| 68 | Trastuzumab | Pemetrexed | Null |
| 69 | Methylseleninic acid | Trastuzumab | Null |
| 70 | Goserelin acetate | Exemestane | (Yang L, Xin T, Rong W U, 2014) |
| 71 | Trabectedin | Tubacin | Null |
| 72 | Toremifene | Tubacin | Null |
| 73 | Toremifene | Trabectedin | Null |
| 74 | Paclitaxel | Capecitabine | (Gradishar *et al.*, 2004) |
| 75 | Sorafenib | Lapatinib | Null |
| 76 | Ixabepilone | Lapatinib | Null |
| 77 | Bevacizumab | Sorafenib | Null |
| 78 | Ixabepilone | Bevacizumab | (Mark Verrill, 2013) |
| 79 | Ixabepilone | Cyclophosphamide | (Yardley *et al.*, 2015) |
| 80 | Cyclophosphamide | Sorafenib | Null |
| 81 | Sorafenib | Docetaxel | Null |
| 82 | Ixabepilone | Docetaxel | Null |
| 83 | Doxorubicin | Gemcitabine | (Pan *et al.*, 2009) |
| 84 | Trastuzumab | Capecitabine | (Bartsch *et al.*, 2007) |

**TABLE III.** Global characteristics of the drug combination network

| Data sets | No. of vertices | No. of edges | No. of non-efficacious drug combinations | Density |
| --- | --- | --- | --- | --- |
| Breast cancer | 41 | 46 | 4 | 0.0561 |
| Colorectal cancer | 18 | 13 | 1 | 0.0850 |
| Lung cancer | 24 | 22 | 2 | 0.0797 |

Breast cancer is the most frequently diagnosed cancer and the leading cause of cancer-related mortality in females worldwide (Zheng *et al.,* 2014), which comprises 22% of all cancers in women (Donahue and Genetos, 2013; Karagoz *et al.,* 2015). We used RWRDC to predict potential efficacious drug combinations. As a result, 100% and 73% of drug-drug pairs which ranked top 5% and 10% in the data set had been validated, respectively. When we took the drug-drug pairs of top 10% the highest predicted efficacious score as the predicted result, all the known efficacious drug combinations contained in the data set would be successfully predicted. Although some predictions have not been validated in databases and literature, there is still a strong possibility that predicted efficacious drug combinations will be available. For example, everolimus-lapatinib was predicted to be 59^th^ by RWRDC and there was no direct evidence that it was an efficacious drug combination. But (Barr *et al.,* 02152016) showed that they were conducting a phase II pilot study, and some HER-2 positive metastatic breast cancer (MBC) were treated with the combination of everolimus and lapatinib for pts. We are confident that the combination of everolimus and lapatinib will perform well in clinical trials. Also, the combination of sorafenib and lapatinib ranked 75^th^ by RWRDC, (Kacan *et al.,* 2014) showed that both sorafenib and lapatinib alone were efficacious in the treatment of breast cancer, also a combination of these two agents may be a promising therapeutic option of the treatment of breast cancer. Besides, sorafenib and docetaxel ranked 81^st^ in the prediction results, although there was no specific literature indicating that they could be combined to treat breast cancer. A phase II trial of sorafenib combined with docetaxel in the treatment of HER2-negative MBC was conducted (Marme *et al.,* 2014). The existing data could not conclude that sorafenib and docetaxel could treat breast cancer, with the development of the trial, there would be clear conclusions soon.

## III. Discussions

The model proposed in this paper is an extension of the classical random walk model in the stochastic process and has a solid mathematical foundation. Mathematically, when the stochastic process reaches a steady-state, we can get the efficacious score *s_ij_*, that is, the possibility of connecting the edges between any vertices *v_i_* and *v_j_*. The drug is treated as a vertex in the modeling process, therefore, there is an edge between the two vertices indicating that the two drugs can be effectively combined, so the model in this paper can be used to predict which drug-drug pairs can be efficacious drug combinations. For a given disease, potential efficacious drug combinations can be predicted. Based on the different types of cross-validation and different data sets, all the results showed that RWRDC had a very strong predictive ability.

Despite the effectiveness of RWRDC, it also has several limitations. Firstly, more types of data can be considered in the model, such as chemical structure similarity between drugs, the semantic similarity between diseases, drug-target interactions, and so on. Introducing more data can improve the prediction accuracy. Furthermore, if the model can quantitatively predict the toxicity and side effects of drug combinations, it will be more convincing. Besides, only the efficacious combination data of two drugs are utilized, and efficacious drug combinations of more than two drugs can be considered in the future. Moreover, we think efficacious drug combinations of more than two drugs can be predicted by a more complex random walk model soon afterward.

In brief, RWRDC is expected to be an important and useful resource for identifying potential efficacious drug combinations and providing useful insights into the underlying mechanisms of drug combinations.

## IV. Conclusion

In this manuscript, we predicted efficacious drug combinations in cancer based on random walk with restart. An efficacious score can be obtained between any two individual drugs by RWRDC. The analysis results suggest that RWRDC is a novel way to discover efficacious drug combinations in cancer, which provides new prospects for cancer treatment. Furthermore, RWRDC is a semisupervised learning framework that can be used to predict combinations of drugs for other complex diseases.

## V. Methods

### A. Data collection and pre-processing

The Drug Combination Database (DCDB) is an available database to collect and organize drug combinations information (Liu *et al.,* 2014), which includes 1363 drug combinations, consisting of 1735 drugs in all. Among these drug combinations, 682 are curated from clinical trials published in the reports, 330 are curated from the FDA orange book and 351 are curated from PubMed articles. There are 1103 efficacious drug combinations in all drug combinations. Besides, DCDB collected 237 non-efficacious drug combinations, which may provide valuable data support for finding efficacious drug combinations. In this paper, we set efficacious drug-drug interactions as positive samples and non-efficacious drug-drug interactions as negative samples.

Because the mortality of breast cancer, colorectal cancer, and lung cancer is very high (Jemal *et al.,* 2010), the drug combinations in DCDB related to these cancers are the data sets studied in this paper. in detail, the data in the database were selected according to the following criteria. iCD10 is an internationally unified disease classification method developed by World Health organization (WHo) (Quan *et al.,* 2005), each drug combination of the DCDB database had at least one ICD10 code. If a drug combination had an ICD10 code: C18, i.e. malignant neoplasm of colon, we recorded this drug combination as colorectal cancer data. Similarly, the drug combinations of C50 and C34 appearing in those ICD10 codes were recorded as breast cancer and lung cancer data, respectively. Also, due to the experimental results of some drug combinations that were still unclear, we did not use those combinations whose effect type was “Need further study” in the DCDB.

We made a simple analysis of the data in the database, as shown in **Table 3**. Here, the edge represented the efficacious drug combination, the vertex represented the drug involved in the efficacious drug combinations. The number of non-efficacious drug combinations was the total number of non-efficacious drug combinations associated with the drugs involved in the efficacious drug combinations. The non-efficacious drug combinations data could be used to verify whether the efficacious score is reasonable (in section B of the Results section). Density referred to the number of edges divided by the number of possible edges, which meant efficacious drug combinations accounted for the proportion of all possible combinations.

**Table 3** shows that the density of efficacious drug combinations used in this paper is very low, that is, only 5% to 9% of all possible drug combinations.

### B. Notations and definitions

Let *G(V,E)* be an undirected graph with the vertex set *V* = {*i*| *i is an individual drug*} and the set of edges *E* = {(*i,j*)}, here *i* and *j* are a pair of efficacious drug combinations that already exist in the current database.

Define the set of positive samples be *PS* = {(*i,j*)|(*i,j*) ∈ E}, the set of candidate samples be *CS* = {(*i,j*)| (*i,j*) ∉ E}, and the set of negative samples *NS* = {(*i,j*)| (*i,j*) *is a non efficacious drug combination*}. In fact, *CS* contains possible *PS* and *NS*.

### C. Random walk with restart algorithm

The random walk on the graph is a transition process by moving from a given vertex to a randomly selected neighboring vertex for each step. We regard the vertex set {*v*_1_, *v*_2_,..., *v_n_*} as a set of states {*s*_1_,*s*_2_,..., *s_n_*}, in a finite Markov chain *ℳ*. The transition probability of *ℳ* is a conditional probability defined as *P(u,v) = Prob(s_t+1_ = v|s_t_ = u)* which means the probability that the *ℳ* will be at *v* at time *t* + 1 given that it was at *u* at time *t*. For any *u* of *V* we have ∑*_v∈V_P*(*u,v*) = 1, we can define a transition matrix *P* ∈ ℝ^|*V*|×|*V*|^ of *ℳ*.

The random walk from *u* to *v* of graph *G* is to choose an edge *e_uv_* randomly. Define transition probability *P(u, v)* as follows:

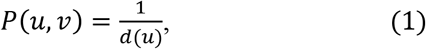

where *d(u)* is the degree of vertex *u*. We denote *D_G_* to be the diagonal matrix containing the vertex degrees of the graph and *A* to be the adjacency matrix of *G*. Thus, *P* can be rewritten in matrix notation as follows:

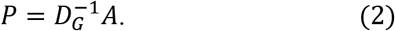

Define *r_t_* ∈ ℝ^|*V*|×1^ as a vector in which the *i*-th element represents the probability of discovering the random walk at vertex *i* at step *t*, then the probability *r*_*t*+1_ can be calculated iteratively by:

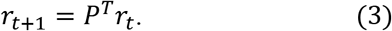

For the random walk with restart (RWR) algorithm (Tong *et al.,* 2006), there is an additional restart item compared to the above algorithm. The probability *r_t+1_* can be calculated iteratively by the following expression:

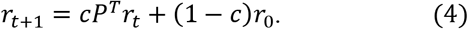

Define initial probability *r_0_ ∈ ℝ^|V|×1^* as a vector in which the *i*-th element is equal to one, while other elements are zeros. And *1 — c* is the restart probability (0 ≤ *c* ≤ 1).

At the beginning of the RWR, we choose a starting vertex *v_i_*, then it would walk based on transition probability. Suppose we come to vertex *v_j_*, then it has a probability of *c* to walk based on transition probability and also have a probability of *1 — c* to restart the walk, that is, to go back to vertex *v_i_*. After some steps, the RWR will be stable, that is, the *L1* norm between *r_t+1_* and *r_t_* is less than an arbitrarily small value. Here we took the cutoff as 10^-6^, and this criterion is often used in some researches (Chen *et al.,* 2012; Chen *et al.,* 2010; Köhler *et al.,* 2008; Li and Patra, 2010). When the RWR is stable, stable probability between vertex *i* and vertex *j* is defined as the *j*-th element of *r_t_* corresponding to the starting vertex is v¿.

### D. Efficacious scoring of drug-drug pairs

Each candidate drug-drug pair can be ranked twice according to RWR. Define 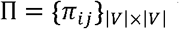 be the stable probability matrix where *π_ij_* indicates the stable probability between vertex *i* and vertex *j*, that is, RWR starts from vertex and the probability of reaching vertex *v_j_* when the process is stable. Briefly speaking, the value of *π_ij_* is the *j-* th element of steady-state probability 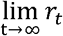 when the *i*-th element of *r_0_* is 1. The significance of *π_ij_* in drug combination can be understood as the possibility of efficacious combination of drug j (i.e. for *n — 1* candidate drugs except for drug i) and given drug i; similarly, *π_ij_* can be understood as the possibility of an efficacious combination of drug i (i.e. for *n — 1* candidate drugs except for drug j) and given drug j. The goal of this paper is to predict the efficacious combination of two drugs, so each drug cannot be combined with itself, but can be combined with other n — *1* drugs. Therefore, define efficacious score *S* = {*s_ij_*}|*V*|×|*V*| as follows:

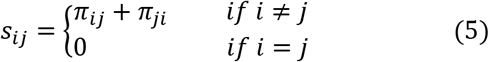

An efficacious score of any two drugs can be calculated by RWRDC. Drug-drug pairs with higher efficacious scores can be expected to have a greater probability to be combinatorial drugs. Experimenters can prioritize the validation of these drug-drug pairs, therefore, the cost to identify potential drug combinations can be significantly reduced. The flowchart of the RWRDC are shown in **Figure 3**.

**Fig. 3.**
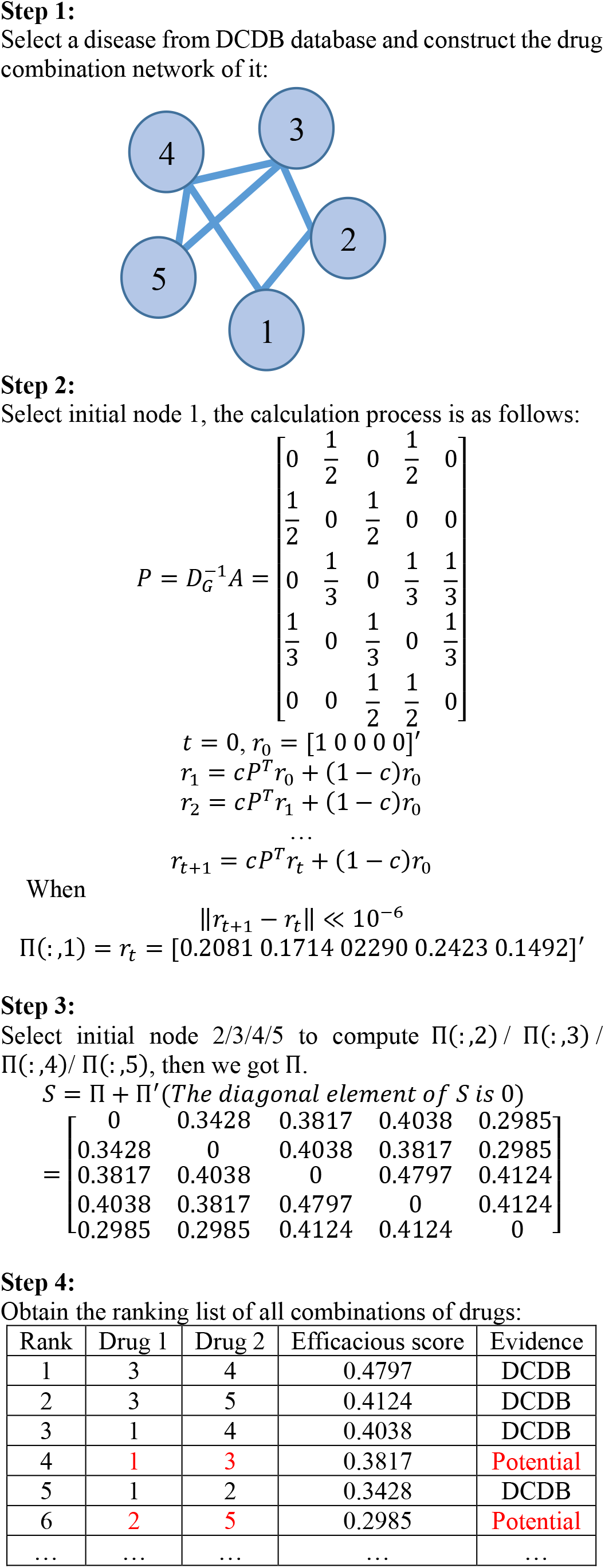
Flowchart of the RWRDC.

The code and dataset of RWRDC are freely available at https://github.com/wangqi27/ERWR

## VI. List of abbreviations

RWRDC: predicting efficacious Drug Combinations in cancer based on Random Walk with Restart
NLLSS: Network-based Laplacian regularized Least square synergistic drug combination prediction
AUROCs: Area Under Receiver Operating Characteristic Curves
ROC: Receiver Operating Characteristic
LOOCV: Leave-One-Out Cross-Validation
DCDB: Drug Combination Database
MBC: Metastatic Breast Cancer
WHO: World Health Organization
RWR: Random Walk with Restart

## VII. Declarations

### Ethics approval and consent to participate

Not applicable.

### Consent to publish

Not applicable.

### Availability of data and materials

Publicly available datasets were analyzed in this study. This data can be found here: http://www.cls.zju.edu.cn/dcdb/. (Temporary DCDB database address: http://public.synergylab.cn/dcdb/.)

The code and dataset of RWRDC are freely available at https://github.com/wangqi27/ERWR.

### Authors’ Contributions

QW conceived the project, developed the prediction method, designed the experiments, implemented the experiments, analyzed the result, and wrote the paper. GY conceived the project, analyzed the result, and revised the paper.

All authors have read and approved the manuscript.

## Acknowledgements

This work was supported by the National Natural science Foundation of China. And the authors would like to acknowledge Dr. Yaqi Zhang for technical support. We thank Dr. Ming Chen *et al.* for making the drug combination data complete, published in ref. (Liu *et al.,* 2014), freely available via the DCDB Database.

## Authors’ Information

**Affiliations**

Academy of Mathematics and systems science, Chinese Academy of Sciences, Beijing, 100190, China.

Qi Wang, Guiying Yan

University of Chinese Academy of sciences, Beijing, 100190, China.

Qi Wang, Guiying Yan

## Corresponding author

Correspondence to Guiying Yan.

